# Tim-3 Promotes Early Differentiation of Tbet^+^ Effector T Cells During Acute Viral Infection

**DOI:** 10.1101/2025.07.08.663709

**Authors:** Priyanka Manandhar, Emily Landy, Kanako Mori, Lawrence P. Kane

## Abstract

The transmembrane protein Tim-3 has received significant attention in recent years as a possible immunotherapy target. This is due to its robust expression on dysfunctional, exhausted, T cells found in the settings of cancer and chronic infection and biochemical evidence suggesting an inhibitory function of Tim-3. However, numerous clinical trials of putative Tim-3 blocking antibodies have yielded modest benefits, at best, in clinical trials for various cancers. Thus, there is a need to more fully understand the function of Tim-3 *in vivo*. Here we have studied the function of Tim-3 early during a T cell response to LCMV Armstrong, which causes an acute viral infection in mice. We show that Tim-3 is rapidly expressed after infection and that the expression of Tim-3 is associated with acquisition of a type I effector phenotype, including expression of T-bet and downregulation of Tcf-1, by both CD4+ and CD8+ T cells. In addition, we demonstrate that knockout or cytoplasmic truncation of Tim-3 attenuates the acquisition of the effector program by T cells after LCMV Armstrong infection. Together, these data help to clarify the role of Tim-3 during acute infection.

## Introduction

A primary function of effector CD8^+^ T cells is to kill cancerous and virally infected cells through the various cytotoxic functions that these cells possess. Naïve CD8^+^ T cells undergo multiple changes in the process of becoming effector cells, one of which is the expression of co-signaling molecules (Jin et al., 2010; John T Chang, 2014; Kaech and Cui, 2012). These molecules have important functions in integrating signals from the TCR, thus regulating T cell function and determining cell fate (Ahn et al., 2018; Chen and Flies, 2013; Kim and Suresh, 2013). T cells also upregulate “checkpoint” inhibitory molecules after activation, to restrain stimulation and limit immunopathology caused by prolonged activation. Some well-known checkpoint inhibitors that are expressed in the context of both chronic infection and cancer include PD-1, CTLA-4, LAG-3 and TIGIT (Anderson et al., 2016; Ferris et al., 2014; Hui et al., 2017). The protein T cell (or transmembrane) immunoglobulin and mucin domain 3 (Tim-3) has also been associated with inhibition of T cells in these settings.

Tim-3, encoded by the *Havcr2* gene, is a type I glycoprotein containing an N-terminal IgV domain, followed by a mucin like domain that has sites for glycosylation, a single transmembrane domain and a C-terminal cytoplasmic domain. Tim-3 is constitutively expressed on multiple immune cell types, including DCs, NK cells, monocytes, macrophages and mast cells and inducibly expressed on T cells (Ferris et al., 2014; Phong et al., 2015). Unlike other checkpoint molecules, Tim-3 does not contain any inhibitory signaling motifs; rather the cytoplasmic domain of Tim-3 contains five tyrosine residues, conserved between mice and humans, which have been shown to be vital for the co-stimulatory activity of Tim-3 (Banerjee and Kane, 2018; Ferris et al., 2014; Lee et al., 2011). Tim-3 can activate signaling pathways like the PI3K and MAPK pathways and is also recruited to the immunological synapse during T cell activation (Avery et al., 2018; Kataoka et al., 2021; Lee et al., 2011). Consistent with these findings, multiple studies have shown that in some contexts Tim-3 may possess co-stimulatory activity in both T cells (Avery et al., 2018; Gorman and Colgan, 2018; Gorman et al., 2014; Kataoka et al., 2021; Lee et al., 2011) and non-T cells like mast cells (Phong et al., 2015).

Naïve T cells undergo many transcriptional, translational and metabolic changes during the process of forming effectors cells (Chung et al., 2021; Kaech and Cui, 2012). These cells are not uniform and can be divided based on the expression of multiple surface markers, effector functions, proliferative capacity and long-term fate (Herndler-Brandstetter et al., 2018; Joshi et al., 2007; Obar and Lefrancois, 2010). Heterogeneity within the CD8^+^ effector T cell population has been defined using markers like KLRG1, CD127, CD27 and transcription factors like T-bet, Eomes, Tcf-1 and Foxo1 (Herndler-Brandstetter et al., 2018; Intlekofer et al., 2005; Joshi and Kaech, 2008; Kaech and Cui, 2012; Obar and Lefrancois, 2010). Tcf-1, another transcription factor closely associated with T cell differentiation, represses Blimp1, which itself is associated with an effector phenotype (Chen et al., 2019; Kallies, 2007; Rutishauser et al., 2009; Tuoqi Wu, 2016; Wu et al., 2015). Additionally, FoxO1 nuclear localization has been directly implicated in the expression of memory genes (Delpoux et al., 2017; Kerdiles et al., 2009; Tejera et al., 2013). These transcription factors in turn are linked to pathways like STAT5 and PI3K signaling (Cannons et al., 2021; Hand et al., 2010; Kim and Suresh, 2013).

Germline deletion of Tim-3 in mice has been shown to increase the frequency of CD8^+^ memory cells at the cost of long-lived effector cells and has also been shown to have a negative effect on the reactivation of memory CD8^+^ T cells (Avery et al., 2018). However, the role of Tim-3 on CD8^+^ T cells at an earlier stage of viral infection remains unknown. Here, we have explored the role of Tim-3 on T cell fate using LCMV Armstrong as a model of acute infection. Our results demonstrate that Tim-3^+^ effector T cells are more terminally differentiated and possess more robust effector function. Using an inducible Tim-3 knockout mouse model and genetically modified T cells that only express a truncated, non-signaling, form of Tim-3, we also show that Tim-3 expression and signaling are required for the formation of a robust effector T cell response.

## Materials and Methods

### Mice

E8i-cre and P14 TCR transgenic mice were obtained from The Jackson Laboratory and bred at the University of Pittsburgh. Tim-3 T2 mutant and *Havcr2^fl/fl^* mice were generated at the University of Pittsburgh Department of Immunology Transgenic and Gene Targeting core. *Havcr2^fl/fl^* mice were generated by using Crispr/Cas9 to insert LoxP sites on either side of exon 4 of the *Havcr2* gene in C57BL/6 zygotes. Disruption of this exon is predicted to result in a premature stop codon 20 bp into exon 5, before the transmembrane domain, leading to nonsense-mediated decay. Insertion of the LoxP sites was initially confirmed by PCR and restriction digest, followed by sequencing of the targeted region. Deletion of the *Havcr2* gene was confirmed at the RNA level.

T2 mutant mice were generated by using Crispr/Cas9 to change the codon for tyrosine 256 in the *Havcr2* gene of C57BL/6 zygotes to a stop codon, resulting in a truncated mutant form of Tim-3. Restriction sites were introduced to facilitate genotyping. When analyzed at baseline, no differences in T cell development were observed in either the thymus or secondary lymphoid organs (unpublished data).

Adoptive transfers of naive P14 cells were performed by injecting 10^4^ CD44^lo^CD62L^hi^ cells retro-orbitally. Both male and female mice 6-8 weeks of age were used for all experiments. Experiments were performed in accordance with protocols approved by the University of Pittsburgh Institutional Animal Care and Use Committee (IACUC).

### Antibodies

The following Abs were used for flow cytometry and flow sorting: CD8α (BioLegend, 563786), TCRβ (BD, 748406), CD4 (BioLegend, 564298), CD45.2 (BD, 564616), CD45.1 (BioLegend, 110724), CD44 (BD, 612799), CD62L (BioLegend, 104438), KLRG1 (Invitrogen, 46-5893-80), CD127 (Tonbo, 60-1271-U100), Tim-3 (BioLegend, 119721 and 119723), T-bet (BioLegend, 644806), Eomes (Invitrogen, 12-4875-80), Tcf-1 (Cell Signaling Technology, 9066S), TCR Va2 (BioLegend, 127809), TCR Vb8.1 (BioLegend, 140103), TNF (BioLegend, 506308), IFN-γ (BioLegend, 505829), biotinylated CD8α (BioLegend, 139360), CD69 (BioLegend, 104536), CD8β (BD, 740278), PD-1 (Invitrogen, 11-9981-81), pERK (Biolegend 675508), pAKT (Cell signaling 4075S), pSTAT5 (BD 562501), pJUN (Cell Signaling 9164) and FoxO1 (Cell Signaling 2880S). The following antibodies were used for *in vitro* blockade: anti Tim-3 RMT-3 (Bioxcell BE0115), anti-Tim-3 5D12 (BioXcell) and anti CD25 antibody (BioXcell BE0012). Representative gating for splenic antigen-experienced T cells is shown in **Fig. S1**.

### LCMV infection

LCMV Armstrong was obtained from Judong Lee and Rafi Ahmed and propagated and tiered as previously described (Welsh and Seedhom, 2008) . Mice were infected with 2x10^5^ PFU of LCMV Armstrong intra-peritoneally or with 4x10^6^ pfu LCMV Clone-13 retro-orbitally. Peripheral blood, spleens or lymph nodes were analyzed by flow cytometry 14 or 35 days post-infection in age and sex-matched groups. Mice were cheek bled using animal lancets. Biotinylated LCMV tetramers were obtained from the NIH Tetramer Core (contract number 75N93020D00005) and used for labelling in RPMI medium with 3% BSA at room temperature for 1 hr. *Ex vivo* peptide restimulation was performed by incubating 2x10^6^ splenocytes in RPMI medium supplemented with 10% BSA, 1% penicillin-streptomycin, 1% L-glutamine, 1% HEPES, 1% nonessential amino acids, 1% sodium pyruvate, and 0.05 mM 2-ME for 5 h in standard tissue culture conditions. A 1:1000 final concentration of brefeldin A containing GolgiPlug was added to the restimulation solution (BD, 55029). GP33, NP396, and GP276 peptides were obtained from Anaspec, and 100 ng/ml pooled peptides were used for *ex vivo* restimulation.

### Flow cytometry, ImageStream analysis and cell sorting

Flow cytometry was performed on a 5-laser Cytek Aurora cytometer and data were analyzed with FlowJo (version 10.6.1).

For ImageStream analysis, spleens were harvested eight days after infection and analyzed by flow cytometry or stimulated with GP33 peptide for the indicated periods of time and co-localization of FoxO1 and DAPI was analyzed on an Amnis ImageStream imaging cytometer.

### xCELLigence cytotoxicity assay

Cytotoxic activity of CD8^+^ T cells was assessed by plating 10^5^ target cells on xCELLigence RTCA plates in c-RPMI media. After culturing target cells for 30 hours, CD44^hi^ Tim-3^hi^ or Tim-3^lo^ cells were sorted and added to wells. Measurements were stopped after 45 hours of co-culture. Normalized cytotoxicity was calculated by dividing final cell growth index by the initial cell growth index.

### RNA and TCR sequencing

C57BL/6 mice were infected with 2x10^5^ pfu of LCMV Armstrong. Spleens were harvested and processed eight days later. Bulk mRNA and TCR sequencing were performed on 10^5^ sorted live CD8^+^CD44^hi^CD62L^lo^ Tim-3^hi^ and Tim-3^lo^ cells. cDNA Library preparation for RNA sequencing was performed using SMART-Seq HT ultra-Low input RNA Kit and for TCR sequencing using the Takara SMARTer mouse TCRα/β profiling kit. DNA libraries were sequenced on the Illumina NextSeq500. mRNA sequencing analysis was performed using Partek Genomics suite, assembled Mus-musculus_mm39 and mapped to Ensembl Transcripts release 106 (Sri Chaparala). TCRseq analysis and visualizations were performed using the R packages Immunarch (V0.6.8) and Mixcr4. RNAseq and TCRseq data are deposited at GEO (record numbers GSE300892, GSE300893 and GSE301002).

### Statistical analyses

Statistical analyses were performed using GraphPad Prism software. Paired and unpaired Student’s t-test and one-way Anova with multiple comparison were used as indicated in figure legends.

## Results

### Tim-3^hi^ CD8^+^ T cells at the effector phase are more activated and terminally differentiated

To assess the phenotype of Tim-3 expressing CD8^+^ effectors during an Armstrong infection, C57BL/6 mice were infected with LCMV Armstrong and splenocytes were analyzed by flow cytometry on day eight after infection. Consistent with previous studies, roughly 30% of CD8^+^ T cells expressed Tim-3 at this time point **(Fig. 1A)**. T cells responding to three LCMV immunodominant peptides (GP33, NP296 and GP376) were quantified using Class I tetramers (**Fig. 1B-G**), which revealed either a small decrease in tetramer^+^ cells among Tim-3^hi^ vs. Tim-3^lo^ cells (GP33, GP276) or no change (NP396), suggesting that Tim-3 promotes the expansion of T cells that do not bind some tetramers, due to either lower affinity or different specificity. Next, we analyzed GP33 tetramer^+^ cells for the activation markers CD44, CD25 and CD69, which were all higher on Tim-3^hi^ cells **(Fig. 1H-J)**. T-bet, a transcription factor associated with CD8^+^ effector differentiation (Joshi et al., 2007) was also significantly higher in these cells (**Fig. 1K**), while conversely the level of Tcf-1 was significantly lower in CD44^hi^CD62L^lo^ antigen-experienced Tim-3^hi^ cells (**Fig. 1L**). Finally, we found that that there was a small, but statistically significant, increase in the proportion of Tim-3^hi^, compared with Tim-3^lo^, cells with a short-lived effector (KLRG1^+^CD127^-^) phenotype (**Fig. 1M**). Taken together, these data suggest that endogenous Tim-3^hi^ cells enter terminal differentiation more quickly, while Tim-3^lo^ cells are still in the process of division and differentiation.

**Fig. 1:**
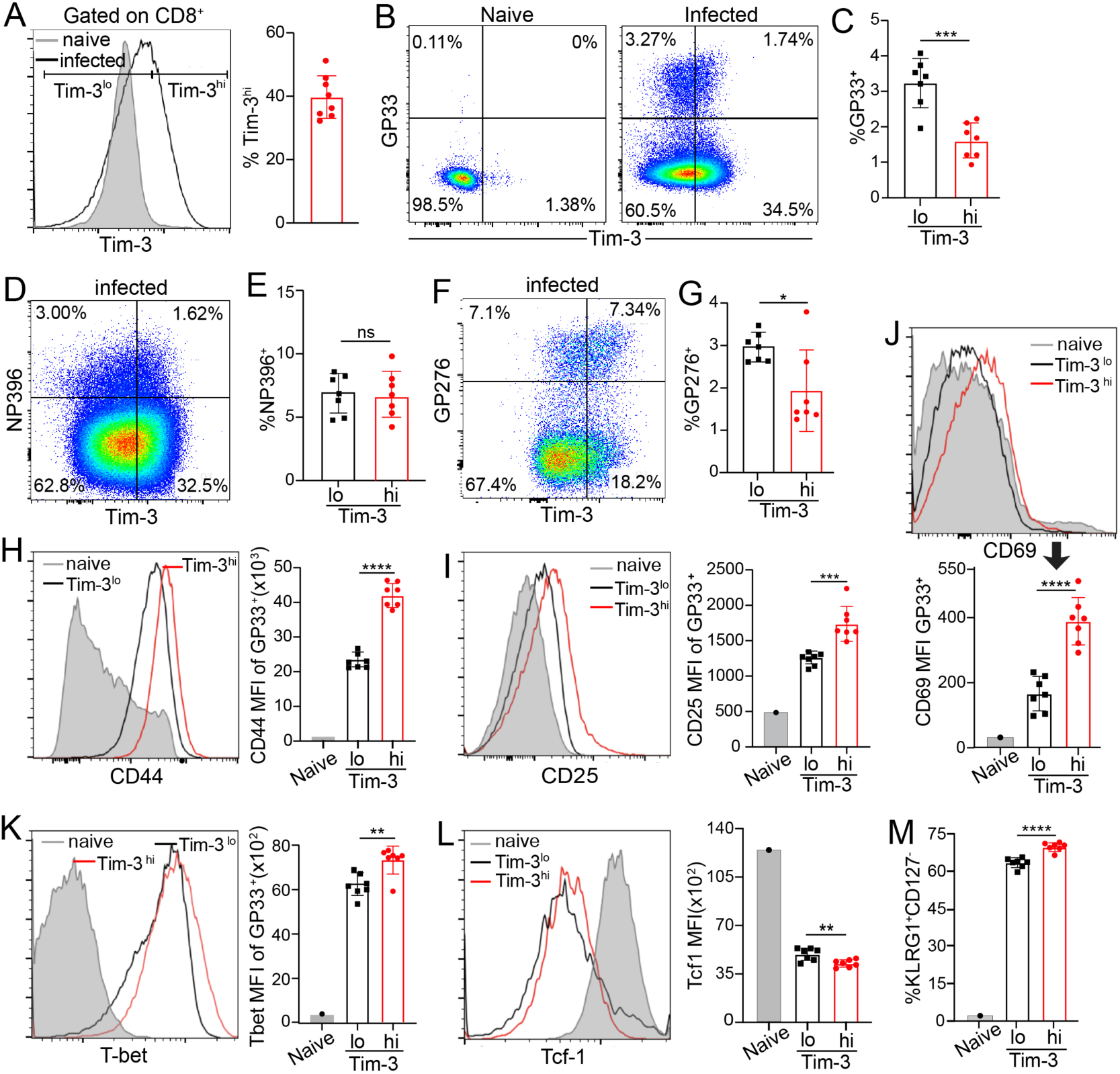
Effector phase Tim-3^hi^ CD8^+^ T cells are more activated and terminally differentiated. C57BL/6 mice were infected with 2x10^5^ pfu of LCMV Armstrong and sacrificed eight days after infection. Spleens were processed and analyzed by flow cytometry. (A) Percent Tim-3^hi^ CD8^+^ T cells. (B-G) GP33^+^, NP396^+^ and GP276^+^ cells of CD44^hi^CD62L^lo^ CD8^+^ T cells. (H-J) CD44, CD25 and CD69 MFI (K) T-bet MFI of GP33^+^ cells and (L-M) Tcf-1 MFI and percentage of KLRG^+^CD127^-^ SLECs of antigen experienced cells. In the graphs, each dot represents a single animal (WT=7). *p<0.05, **p < 0.01 and ****p<0.0001. Data are representative of two experiments.

### Tim-3^hi^ CD8^+^ T cells undergo more rapid differentiation

While adaptive immunity and an effective T cell response take at least a week to develop, LCMV Armstrong is already cleared after one week of infection, while viral titers are highest around day three to four. To understand the dynamics of Tim-3^hi^ vs. Tim-3^lo^ CD8^+^ T cells, we focused earlier after infection, on day four, a time point when LCMV can still be detected. We found that a higher percentage of antigen experienced CD8^+^ T cells expressed Tim-3 at this time point, compared with day eight. Thus, 50-60% of antigen-experienced CD8^+^ T cells were Tim-3^+^ positive at day four after infection **(Fig. 2A)**. Similar to day eight after infection, Tim-3^+^ cells also expressed higher levels of CD44 and CD25 **(Fig. 2B-C),** indicating that Tim-3 marks a subset of CD8^+^ effectors that are activated very early after infection. The hallmark effector markers KLRG-1 and T-bet were also significantly higher in Tim-3^+^ cells (**Fig. 2D-E**), suggesting that these cells may be preferentially forming terminal effectors, as opposed to memory precursors, very early in the infection. Other checkpoint molecules such as PD-1, LAG-3 and TIGIT are also known to be expressed during the early stages of an LCMV infection and Tox, a regulator of exhaustion that is also expressed during acute stimulation, was also significantly higher in Tim-3^+^ cells **(Fig. 2F)**.

**Fig. 2:**
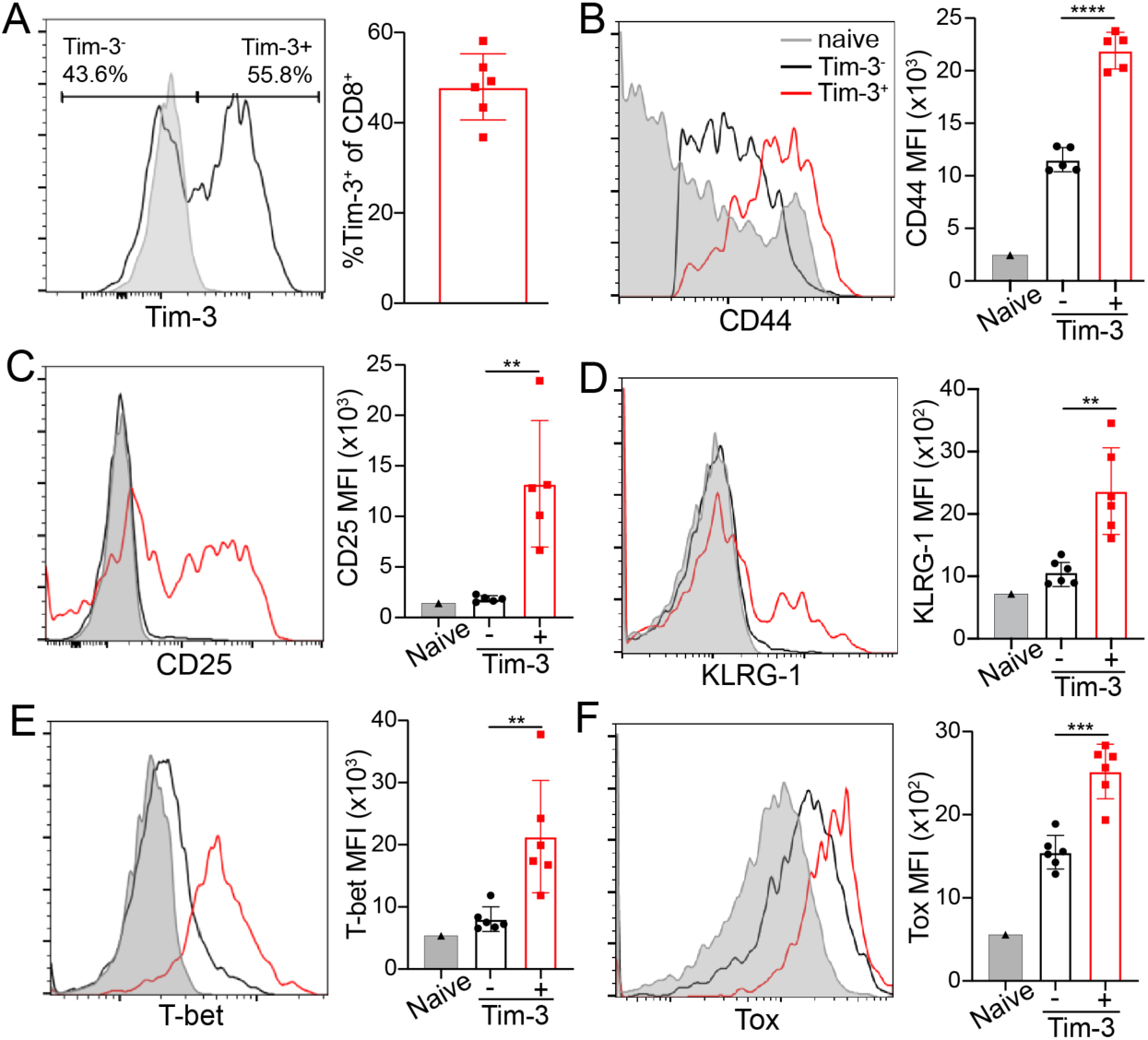
Tim-3^+^ CD8^+^ T cells undergo more rapid differentiation. Mice were infected with 2x10^5^ pfu of LCMV Armstrong and sacrificed four days after infection. Spleens were processed and analyzed by flow cytometry. (A) Percent Tim-3^+^ of CD8^+^ T cells (B-E) CD44, CD25 and KLRG-1 MFI and (E-G) T-bet and Tox MFI of CD44^hi^CD62L^lo^ CD8^+^ T cells. Statistical significance was derived from Student’s t-test. In the graphs, each dot represents a single animal (WT=5). \*\**p* < 0.01, ***p<0.001, ****p<0.0001. Data are representative of two experiments.

### Tim-3^hi^ CD8^+^ effectors have enhanced cytokine production and cytotoxic function

Since endogenous Tim-3^hi^ CD8^+^ T cells were more activated and terminally differentiated in comparison to Tim-3^lo^ cells, the functional capabilities of these cells were assessed by measuring cytokine production after LCMV peptide re-stimulation. A characteristic of exhausted Tim-3 expressing CD8^+^ T cells in a setting of chronic stimulation is that these cells are “dysfunctional” and unable to produce cytokines, a key function of CD8^+^ T cells in the process of viral clearance (Ferris et al., 2014; Wherry et al., 2007). To measure cytokine production, splenocytes from C57BL/6 mice at day eight after infection were stimulated with pooled GP33, NP396 and GP276 peptides plus brefeldin A for five hours and intracellularly labelled for cytokines and analyzed by flow cytometry. Tim-3^hi^ CD8^+^ effector T cells gated on CD44^hi^ cells had significantly higher levels of IFN-γ, TNF and IFN-γ/TNF “double producers” **(Fig. 3A-D)**.

**Fig. 3:**
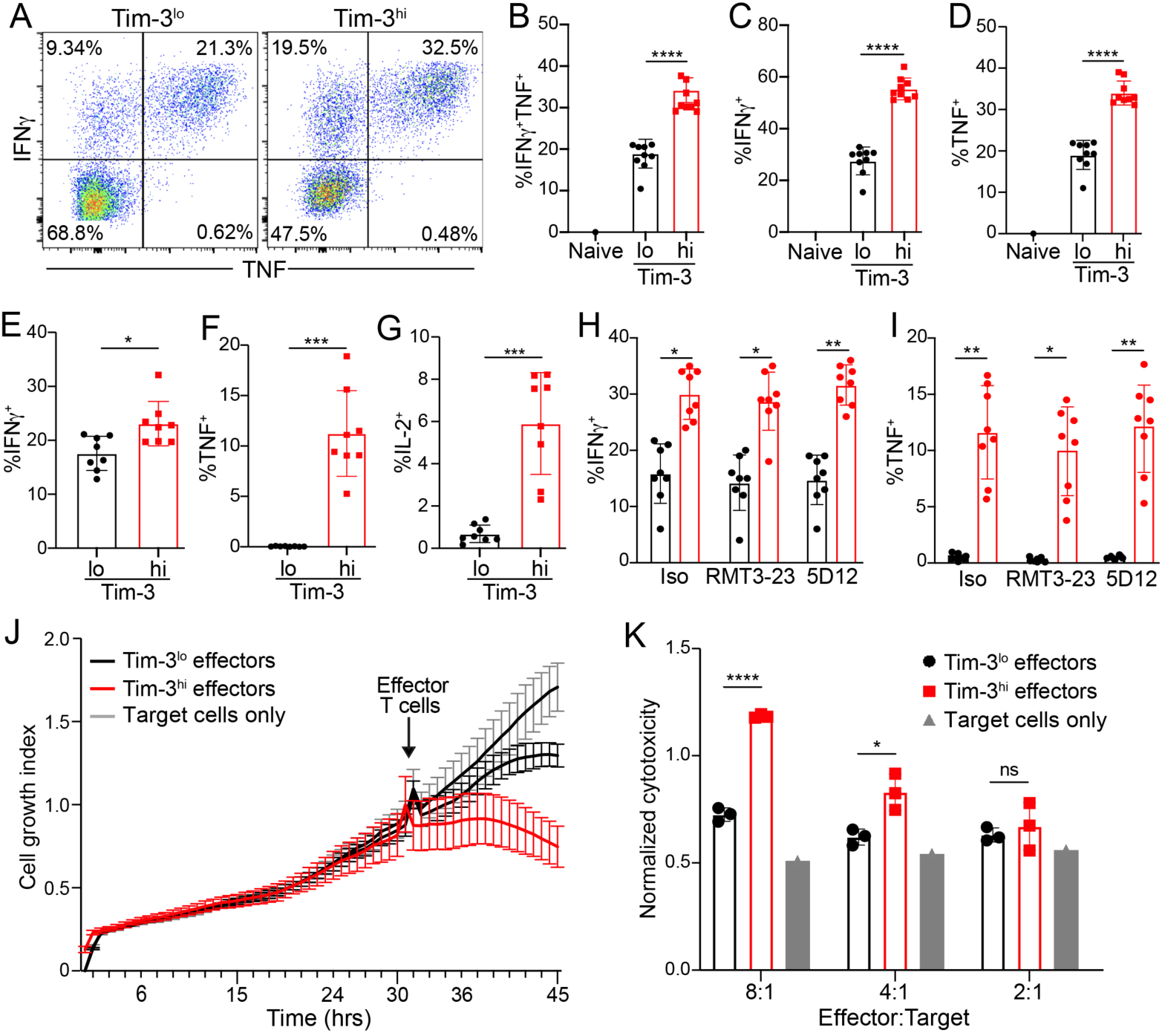
Tim-3^hi^ CD8^+^ effector T cells have enhanced cytokine production and cytotoxic ability. C57BL/6 mice were infected with 2x10^5^ pfu of LCMV Armstrong and sacrificed eight days after infection. Spleens were stimulated with pooled LCMV peptides or anti-CD3 antibody with brefeldin A for five hours. Xcelligence assay was performed using MC38-GP and Tim-3^lo^ and Tim-3^hi^ CD8^+^ effector T cells sorted from day 8 LCMV-infected mice. (A-F) cytokine production after five hours of peptide restimulation. (G-I) Cytokine production after stimulation with anti-CD3 antibody. (J-L) cytokine production after peptide restimulation and blockade with indicated isotype control or Tim-3 antibodies. (M-N) xCELLigence cytotoxicity (cell growth index curve and normalized cytotoxicity). In panels A-L, each dot represents an individual mouse (n=9); in panels M-N, each dot represents a technical replicate. Statistical significance was derived from Student’s t-test. (WT=7; KO=5). *p<0.05, **p < 0.01, ****p<0.0001. Data are representative of two experiments.

Production of the cytokines IFN-γ, TNF and IL-2 were also significantly higher when splenocytes were stimulated with anti-CD3 mAb alone, suggesting that Tim-3^hi^ cells can produce cytokines without anti-CD28 co-stimulation **(Fig. 3E-G)**. Multiple ligands for Tim-3 have been discovered, which have been reported to function as both positive and negative regulators of Tim-3, in different contexts (Dardalhon et al., 2010; Huang, 2015; Smith et al., 2021; Wolf et al., 2020). To assess the role of ligand binding in the functional properties of Tim-3^hi^ CD8^+^ effectors, RMT3 and 5D12 were used to block Tim-3 in splenocytes during re-stimulation. Thus, addition of either of these antibodies did not affect cytokine production, suggesting that ligand binding did not play a major role in the functional activity of Tim-3^hi^ CD8^+^ T cells **(Fig. 3H-I)**.

Finally, the cytotoxic capability of Tim-3^hi^ vs Tim-3^lo^ cells was measured using MC38 tumor cells expressing the glycoprotein (GP) of LCMV as target cells. The cell growth index of target cells that were co-cultured with Tim-3^hi^ target cells was significantly lower than target cells co-cultured with Tim-3^lo^ cells **(Fig. 3J)**, indicating greater cytotoxic activity of the Tim-3^hi^ cells. Normalized cytotoxicity by Tim-3^hi^ cells was also significantly higher than that mediated by Tim-3^lo^ cells and was dose-dependent **(Fig. 3K)**. These findings further establish that Tim-3^hi^ CD8^+^ effector T cells are more-differentiated and have superior cytotoxic activity on a per-cell basis.

### Upregulation of distinct effector genes and focusing of the TCR repertoire of Tim-3^hi^ CD8^+^ effector T cells

To further study the effects of Tim-3 expression on CD8^+^ T cells, antigen-experienced CD44^hi^CD62L^lo^ Tim-3^lo/hi^ cells from LCMV Armstrong-infected mice were sorted and analyzed by RNA sequencing. Tim-3^lo^ and Tim-3^hi^ cells showed distinct clustering by PCA analysis **(Fig. S2A)**. A volcano plot of genes significantly upregulated or downregulated at least two-fold reveals that there are more genes upregulated in the Tim-3^hi^ cells (**Fig. S2B**), consistent with their more-differentiated status. GSEA analysis of genes differentially expressed in the Tim-3^hi^ vs. Tim-3^lo^ cells revealed multiple pathways with distinct positive correlations (**Fig. S2C-D**). These pathways included apoptosis and programmed cell death, cell division, DNA replication and mitotic cell cycle. Metabolic pathways were also positively coregulated and other miscellaneous cell process pathways included regulation of translation, DNA repair and cell growth pathways, consistent with more efficient activation and effector differentiation of Tim-3^hi^ CD8^+^ T cells. Specific genes of interest upregulated by a fold change of two or greater in Tim-3^hi^ cells included *Lag3*, *Pdcd1*, *Ezh2* and *Gzmb* (**Fig. S2E**). Other effector associated genes that were upregulated included *Il2ra* and *Tox*, while a variety of naïve or memory associated genes were downregulated in Tim-3^hi^ cells including *Tcf7*, *Bach2*, *Cxcr5*, *Ccr7*, *Il7r*, *Klf2*, *Sell* and *Bcl6*. Bcl-2 family members which are primarily associated with a memory or naïve phenotype were also significantly lower in Tim-3^hi^ cells, while *Tox* was higher at the transcript level, consistent with flow cytometry analysis shown above. Memory-associated cell-surface markers like *Il7ra*, *Ccr7* and *Sell* were also significantly lower in Tim-3^hi^ cells, while activation-associated surface markers *Pdcd1*, *Lag3* and *Il2ra* were higher. Cytokines and effector molecules like *Ifng*, *Il18*, *Gzmb* and *Gzmc* were also significantly higher in Tim-3^hi^ CD8^+^ effectors, together showing that Tim-3^hi^ cells have an overall more effector-like phenotype **(Fig. S2E)**.

We next considered the role of Tim-3 in shaping the repertoire of responding CD8^+^ T cells. TCR sequencing revealed that the Tim-3^hi^ CD8^+^ population had significantly lower *TCRa* and *TCRb* diversity, as determined by the Chao1 diversity estimator **(Fig. S3A-B)**. While there was sharing of clonotypes between the two populations among the top 30 clones, these clones had occupied a larger proportion in Tim-3^hi^ cells compared to Tim-3^lo^ cells **(Fig. S3C)**. In terms of the *TCRb* clones, the top 10 and 100 clones also occupied more repertoire space compared to Tim-3^lo^ effectors **(Fig. S3D)**. Finally, we also assessed *TCRa* and *TCRb* diversity using the Hill diversity number. Again, the Tim-3^hi^ cells displayed reduced diversity of both chains (**Fig.**

**S3E-F**). Taken together, these data demonstrate that a smaller number of Tim-3^hi^ TCR clones occupy a larger space in the TCR repertoire, suggesting that Tim-3 plays a role in focusing of the T cell response.

### Tim-3^hi^ cells are elevated in multiple pathways and exhibit an effector-like phenotype associated with inhibition of FoxO1 nuclear entry

The co-stimulatory function of Tim-3 has previously been shown to require MAPK and mTOR-Akt signaling (Avery et al., 2018; Kataoka et al., 2021; Lee et al., 2011). FoxO1 is a transcription factor that is downstream of the Akt pathway. Once phosphorylated by Akt, FoxO1 is sequestered outside the nucleus, which prevents it from driving transcription of quiescence-associated target genes (Delpoux et al., 2017; Delpoux et al., 2021; Hedrick, 2012; Zhang et al., 2016). Tim-3^hi^ CD8^+^ effectors also had significantly higher pERK and pAkt^473^ levels **(Fig. 4A-B)**, consistent with previous studies from our group and others (Avery et al., 2018; Lee et al., 2011; Smith et al., 2021). There was also a higher level of JNK activation in these cells, as shown by pJun staining, while pSTAT5 staining was not statistically significant between the populations (**Fig. 4C-D**). FoxO1 nuclear localization was also quantified using the Amnis ImageStream system to measure the nuclear localization of FoxO1 after stimulation with GP33 peptide. Thus, nuclear localization of FoxO1 was significantly lower in Tim-3^hi^ cells compared to Tim-3^lo^ cells after fifteen minutes of stimulation, while total FoxO1 levels were not significantly different **(Fig. 4E-H)**. Representative FoxO1 staining is shown in **Fig. 4I**. Together, these data are consistent with the association between Tim-3 expression, signaling through MAPK and Akt and exit from quiescence, through inhibition of FoxO1 activity.

**Fig. 4:**
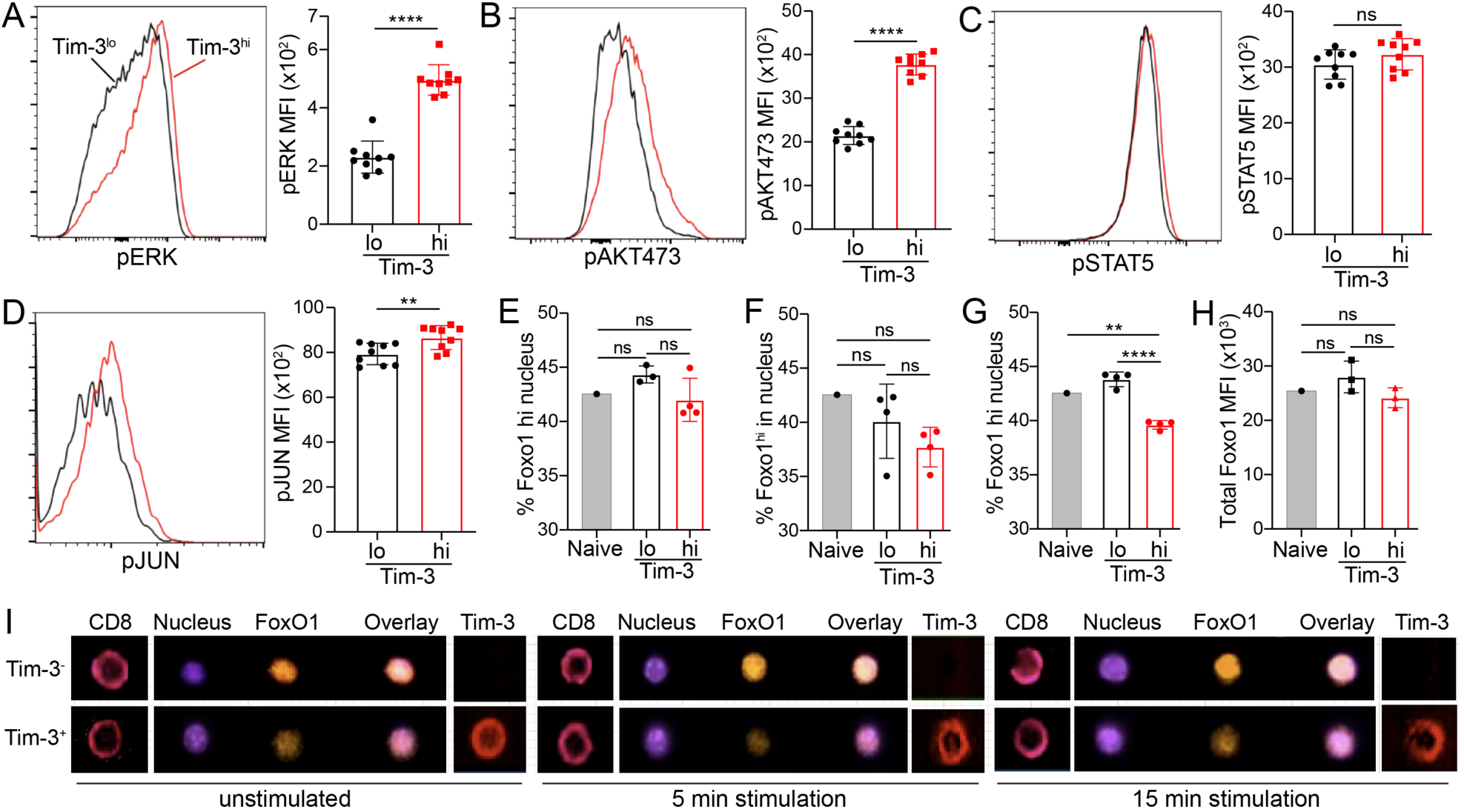
Multiple signaling pathways are upregulated and FoxO1 nuclear localization is inhibited in Tim-3^hi^ effector CD8^+^ T cells. C57BL/6 mice were infected with 2x10^5^ pfu of LCMV Armstrong virus and spleens were harvested at day 8. Splenocytes were stained *ex-vivo* for flow cytometry or stimulated with GP33 peptide and analyzed by ImageStream. pERK, pJUN and pAKT^473^ MFI in CD44^hi^CD62L^lo^ CD8^+^ T cells. (E-G) Proportion of nuclear FoxO1 directly *ex vivo* (E) or after peptide stimulation for 5 minutes (F) or 15 minutes (G). (H) Total FoxO1 MFI *ex vivo*. (I) Representative images of CD8, DAPI, Foxo1 and Tim-3 from ImageStream analysis. Statistical significance was derived from Student’s t-test (A-C) or one-way Anova with Tukey’s multiple comparison (D-G). In the graphs, each dot represents a single animal (n=9 or n=4). Approximately 3000-5000 cells were acquired per animal in (D-G). \*\**p* < 0.01, *\*\*\*\*p<*0.0001. Data are representative of two experiments.

### Tim-3 is required for a robust CD8^+^ effector response to LCMV Armstrong

To assess the function of Tim-3 during the effector stage of acute infection, Tim-3 floxed mice were crossed to E8i-Cre mice (Tim-3^CD8^ KO). These mice, and Cre-only controls, were infected with 2x10^5^ pfu of LCMV Armstrong and splenocytes were analyzed on day eight, as in the studies described above. First, we validated E8i-driven Tim-3 KO, which we found to be extremely efficient (**Fig. 5A**). Consistent with a possible role for Tim-3 in activation and proliferation, as discussed above, the total number of splenocytes was significantly lower in KO mice **(Fig. 5B)**. In addition, the total number of CD8^+^ T cells was slightly lower in the KO **(Fig. 5C)**. While the total number of GP33^+^ cells was slightly (but not significantly) lower, the number of NP396^+^ and GP276^+^ cells was significantly lower in the KO (**Fig. 5D-F**). Viral titers (measured by qPCR) from the spleen (**Fig. 5G**) and the liver (**Fig. 5H**) were not statistically significant between WT and CD8-specific Tim-3 KO mice, consistent with the robust CD8 effector response in C57BL/6 mice. We next assessed markers of T cell activation and differentiation. Thus, expression of the transcription factor Tcf-1 was higher in the KO in GP33^+^ and GP276^+^ CD8^+^ effectors **(Fig. 5I)**, suggesting the KO cells were less-differentiated. Consistent with this, the terminal effector differentiation markers Klrg1 and T-bet were significantly lower in the Tim-3 KO cells **(Fig. 5J-K)** indicating that Tim-3 supports an effector like phenotype during CD8^+^ T cell activation and differentiation. Furthermore, Tim-3 also appeared to be required for the full functionality of effector CD8^+^ T cells, as the level of granzyme B after peptide stimulation was lower in Tim-3 KO T cells **(Fig. 5L)**.

**Fig. 5:**
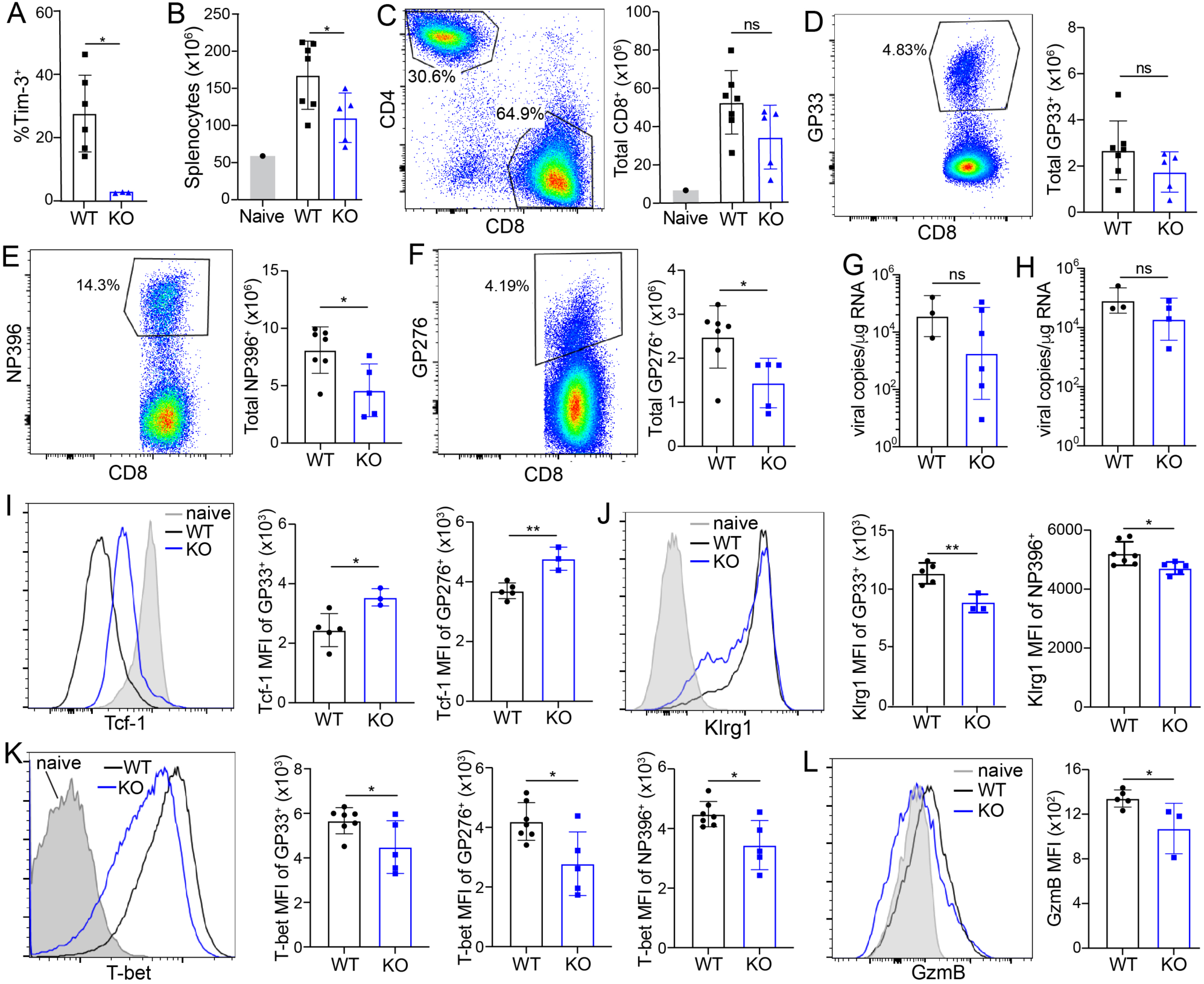
Tim-3 is required for a robust CD8^+^ effector T cell response to LCMV Armstrong. E8i-Cre or *Havcr2^fl/fl^* E8i-Cre (KO) mice were infected with 2x10^5^ pfu of LCMV Armstrong virus and spleens were harvested at day 8. (A) Tim-3 expression on CD8^+^ T cells. (B) Total splenocytes. (C) Total CD8^+^ T cells. (D-F) Percent tetramer^+^ cells for GP33 (D), NP396 (E) and GP276 (F). (G-H) Viral titers by qPCR at day 4 after infection. (I-K) Expression of markers of differentiation (I – Tcf-1; J – Klrg1; K – T-bet) on antigen-experienced T cells stained with the indicated tetramers. (L) GzmB MFI in CD44^+^ cells after 5 hours of peptide re-stimulation.

Tim-3 was first described as a relatively specific marker for Th1 T cells, as compared with naïve and Th2 T cells, which do not express any detectable Tim-3 protein (Monney et al., 2002). Therefore, we also bred the floxed Tim-3 mice to mice expressing CD4-driven Cre (Lee et al., 2001), to drive deletion of Tim-3 from all α/β T cells. Using these mice, we focused on the effects of Tim-3 deficiency on the activation and differentiation of CD4^+^ T cells in response to infection with LCMV Armstrong, again around the peak of the T cell response to infection. While the total number of CD4^+^ T cells in the spleens of infected mice was not different (**Fig. 6A**), there was, as expected, a substantial decrease in the proportion of Tim-3^+^ CD4^+^ T cells in the Tim-3^CD4^ KO animals (**Fig. 6B**). We next examined CD4^+^ T cells specific for the GP66 epitope from LCMV and found that there were comparable numbers of such cells in WT and KO animals (**Fig. 6C**). However, as with CD8^+^ T cells, there were phenotypic differences in the KO CD4^+^ T cells, with Tim-3 KO CD4^+^ T cells displaying moderately lower levels of the receptor KLRG1 (**Fig. 6D**) and the Th1 signature transcription factor T-bet (**Fig. 6E**).

**Fig. 6:**
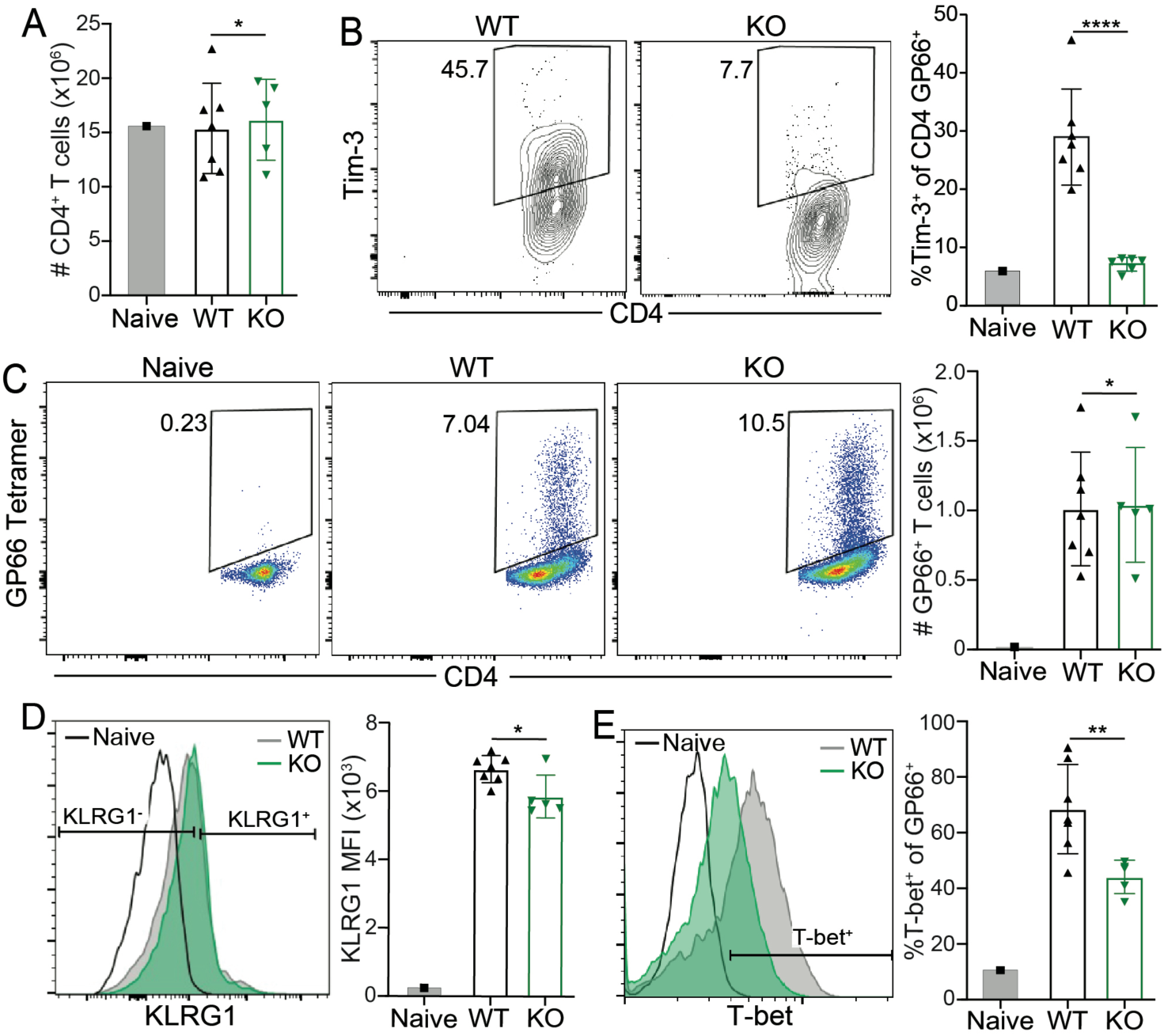
Tim-3 is required for optimal differentiation of effector Th1 T cells at the peak of the response to LCMV Armstrong. Mice with CD4-driven Cre alone (WT) or together with *Havcr2^fl/fl^* (KO) were infected i.p. with LCMV Armstrong. On day 7, splenic CD4^+^ T cells were analyzed by flow cytometry. (A) Total number of splenic CD4+ T cells. (B) Validation of Tim-3 KO efficiency in CD4^+^ T cells. (C) Quantitation of LCMV-specific CD4^+^ T cells based on staining with GP66 tetramers. (D-E) Phenotype of CD4^+^ T cells in WT and Tim-3 KO CD4^+^ T cells, based on staining for KLRG1 (D) and T-bet (E).

To study the role of Tim-3 on CD8^+^ T cells in the context of a fixed TCR, Tim-3 KO mice were crossed to LCMV GP33-specific P14 TCR Tg mice. Additionally, to study the role of signaling via the intracellular domain of Tim-3, mice expressing a germline mutant form of Tim-3 lacking the cytoplasmic domain (T2 mutant) were also crossed to P14 TCR Tg mice. The experimental design and schematic of the T2 allele are shown in **Fig. 7A-B**. Briefly, P14 T cells (10^3^) were adoptively transferred to congenically marked mice and infected with LCMV Armstrong. At eight days after infection, P14 cells from the spleen were analyzed by flow cytometry and bulk RNA sequencing. Although total P14 cell numbers were not significantly different (data not shown), T-bet was significantly lower in P14 (KO and T2) cells as was the transcription factor Blimp-1 **(Fig. 7C-D)**. Bulk RNA sequencing of P14 cells revealed that the WT and KO cells segregated independently by PCA analysis, as expected. However, the T2 mutant CD8^+^ T cells did not cluster directly with the KO cells but rather displayed an apparently intermediate phenotype **(Fig. 7E)**. This suggests that there may be some function of Tim-3 in this context that is cell-extrinsic in nature. Nonetheless, GSEA analysis showed that transcripts from both the T2 mutant and Tim-3 KO P14 cells were negatively correlated with a dataset that compared effector vs. memory genes, indicating that in the absence of Tim-3 signaling, effector CD8^+^ T cells showed less of an effector signature **(Fig. F-G)**.

**Fig. 7:**
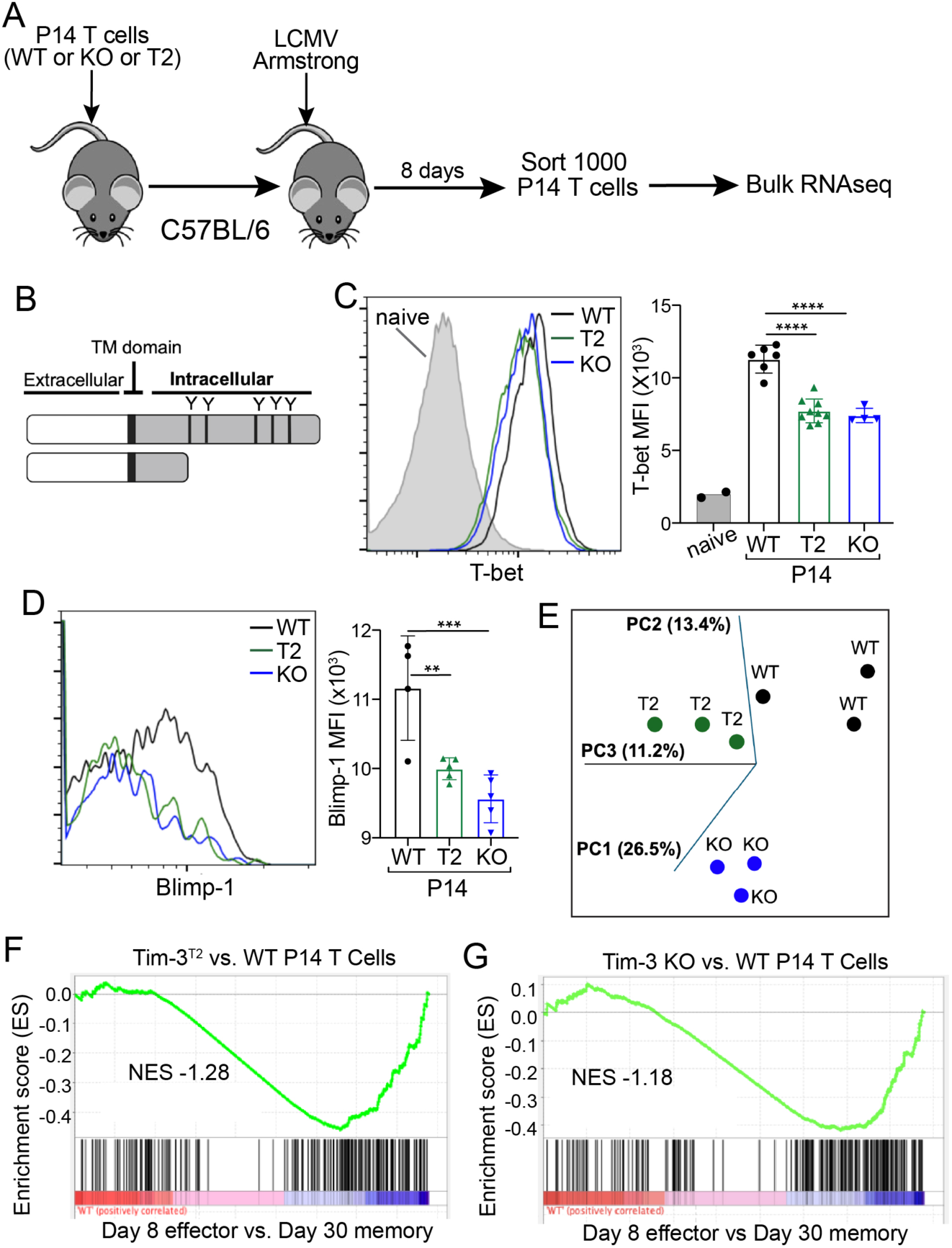
Role of Tim-3 signaling in CD8^+^ T cell phenotype and genotype during acute LCMV infection. (A) Naïve WT or *Havcr2^fl/fl^* E8i-Cre P14 T cells (10^3^) were adoptively transferred into congenically marked mice and infected with LCMV Armstrong. (B) Schematic of the domain structure of WT and T2 forms of Tim-3. (C-D) P14 cells were analyzed by flow cytometry for expression of T-bet (C) or Blimp1 (D). (E-F) P14 cells were sorted and bulk RNA sequencing was performed. (E) Three-dimensional PCA analysis of the transcriptome of WT, T2 and Tim-3 KO P14 cells. (F-G) GSEA analysis of genes differentially expressed in T2 (F) or KO (G) vs. WT P14 CD8^+^ T cells, compared to day 8 effector vs. day 30 memory cells from the dataset of Kaech and colleagues.

## Discussion

Tim-3 is primarily known as a terminal exhaustion marker expressed on dysfunctional T cells, but its expression has also been reported on activated T cells during acute infection. The role of Tim-3 on CD8^+^ T cells during the effector response to an acute viral infection has, to our knowledge, not been dissected in detail. Our data show that Tim-3 marks a highly differentiated and activated population of CD8^+^ T cells during LCMV Armstrong infection. Mature Tim-3^+^ CD8^+^ effectors had a distinctly differentiated effector phenotype at day eight after infection.

Additionally, Tim-3 upregulation was also associated with an effector phenotype as early as four days after infection. Activation markers CD44, CD25 and CD69 were upregulated on Tim-3^+^ effectors, along with T-bet, a transcription factor well-known to be associated with an effector phenotype and which is upregulated by TCR stimulation. LCMV-specific T cells staining with tetramers containing immunodominant epitopes (GP33 and GP276) were found in significantly lower numbers in the absence of Tim-3.

We previously reported that CD8^+^ T cells ectopically expressing Tim-3 displayed enhanced *in vitro* proliferation after two days of TCR stimulation but had lower proliferation by four days (Kataoka et al., 2021). Taken together with the *in vitro* proliferation assay, this suggests that Tim-3^hi^ CD8^+^ T cells have reached differentiation faster and are no longer proliferating. Signaling pathways known to lie downstream of TCR were also lower in Tim-3^lo^ cells *ex vivo* and these cells had a correspondingly higher level of nuclear localization of the transcription factor FoxO1. Active FoxO1 in the nucleus has previously been shown to repress T-bet mediated effector functions in CD8^+^ T cells (Rao et al., 2012).

A hallmark of mature, differentiated cytotoxic CD8^+^ T cells is the production of cytokines such as granzymes, perforin, IFN-γ, TNF and IL-2 (Kaech and Cui, 2012). Tim-3 also marked a population of effector CD8^+^ T cells that are functionally superior, as shown by cytokine production after LCMV peptide or anti-CD3. Furthermore, Tim-3^hi^ cells also showed higher cytotoxicity in the *ex-vivo* killing assay using the LCMV GP-expressing MC38 cell line. Binding of the phosphatidylserine has been shown to support the co-stimulatory function of Tim-3; hence we utilized anti Tim-3 antibodies known to block the binding of phosphatidylserine (Sabatos-Peyton et al., 2018; Smith et al., 2021). However, these antibodies did not affect, either positively or negatively, the elevated cytokine production by Tim-3^hi^ cells. The activated and terminally differentiated terminal phenotype of Tim-3^hi^ CD8^+^ effectors was confirmed and extended using bulk RNA sequencing of antigen-experienced CD8^+^ T cells. Interestingly, the TCR clonality of Tim-3^hi^ effectors appeared to be lower in terms of both TCRα and TCRβ, compared with activated Tim-3^lo^ cells. In other words, the most abundant clones appeared to occupy more space in the TCR repertoire of Tim-3^hi^ cells. Overall, this suggests that clones that expressed Tim-3 may have expanded faster during the initial stages of proliferation and reached a terminal effector phenotype faster, since focusing of the TCR repertoire is indicative of a more mature response (Zehn et al., 2009).

To elucidate the functional role of Tim-3 signaling in a CD8 response during an acute viral infection, we generated a CD8-specific KO model of Tim-3 by crossing our previously described floxed Tim-3 mice (Avery et al., 2018) to mice carrying the E8i-Cre allele (Maekawa et al., 2008). Infecting these mice with LCMV Arm revealed that lack of Tim-3 expression led to a reduction in effector CD8^+^ T cells between days four and eight after infection, an effect that was particularly striking at the later time points. Tim-3 KO mice had a significantly lower number of splenocytes and the total number of CD8^+^ T cells trended lower, although the latter was not statistically significant. LCMV tetramer NP396^+^ and GP33^+^ cells were significantly lower on KO mice at day eight of infection. Viral titers were measured by qPCR after four days of infection from the spleen and liver but were not found to be significantly different. Any differences in viral titers present at this point may have been obscured by the high titers of virus seen at this time point (Matloubian et al., 1994). Additionally, expression levels of other hallmark effector markers like Klrg1 and T-bet were significantly lower in KO mice. Restimulation of KO T cells with pooled LCMV peptides also revealed a reduction in the level of granzyme B protein. Taken together, these data highlight the role of Tim-3 in the formation of a robust CD8^+^ T cell response We also studied the effects of Tim-3 on CD8^+^ T cell fate using LCMV GP33-specific TCR transgenic P14 mice, taking advantage of mice expressing a non-signaling mutant of Tim-3, the T2 mutant, which lacks the cytoplasmic tyrosine residues (Manandhar et al., 2024). In earlier work, we showed that Tim-3 mutants lacking the tyrosine residues have defective co-stimulatory signaling in T cell lines (Lee et al., 2011). Thus, significant differences in the effector associated transcription factors T-bet and Blimp-1 were seen in effector P14 T cells lacking either Tim-3 expression or its signaling domain. P14 cell numbers were not significantly different, possibly due to high TCR levels on P14 cells. Bulk RNA sequencing analysis of effector P14 cells also indicated that Tim-3 KO P14 T cells cluster independently of WT cells, while Tim-3^T2^ P14 T cells had an intermediate phenotype, which may be due to ligand interaction of Tim-3. However, GSEA analysis revealed that both KO and T2 mutant T cell transcriptomes negative correlated with a day eight vs day 30 memory gene set, compared with WT T cells.

Although it marks terminally exhausted CD8^+^ T cells in chronic infections and tumors, Tim-3 distinctly lacks inhibitory or switch motifs typically associated with recruitment of phosphatases (Banerjee and Kane, 2018; Ferris et al., 2014). Ligands known to modulate Tim-3 signaling include galectin-9, HMGB-1, Ceacam-1 and phosphatidylserine (Banerjee and Kane, 2018; C M. Smith*, 2021; Ferris et al., 2014). While the interaction of Tim-3 with galectin-9 can induce apoptosis in Th1 cells (Zhu et al., 2005), phosphatidylserine plays a role in the clearing of apoptotic bodies through binding to Tim-3 (DeKruyff et al., 2010). HMGB1 interaction with Tim-3 has been shown to dampen the innate immune response to tumor-associated nucleic acids (Chiba et al., 2012). Blockade of Tim-3 has yielded variable results in different mouse models, but dual blockade of Tim-3 and PD-1 has been shown to be more effective in reducing tumor burden and controlling chronic LCMV infection, compared with blockade of PD-1 alone (Ahn et al., 2018; Sakuishi et al., 2010). Tyrosine residues present in the intracellular domain of Tim-3 appear to play a critical role in Tim-3 signaling, as demonstrated through both *in vitro* and *in vivo* models (Kataoka et al., 2021; Lee et al., 2011). The T2 mutation showed a loss of Tim-3 mediated co-stimulation in *in vitro* assays (Lee et al., 2011), as well as impaired effector differentiation of P14 cells in the setting of LCMV Armstrong infection. In the present study, a more quiescent or memory-like phenotype was observed in T2 mutant T cells, as shown by RNA sequencing, indicating that Tim-3 signaling can promote effector differentiation of CD8^+^ T cells.

Recent studies have revealed diverse subsets of CD8⁺ T cells in both chronic and acute Armstrong infections, distinct from terminally exhausted or memory exhausted cells. T precursor exhausted cells (Tpex) have been detected at early time points during LCMV Armstrong infection and conversely, stem-like T cells expressing memory markers have been detected in early chronic LCMV Cl-13 infections (Chu et al., 2025; McManus et al., 2025). These findings suggest that a variety of T cell populations are formed early during infections and serve various purposes during an anti-viral immune response. Thus, Tpex cells expressed exhaustion-associated genes such as *Tox, Lag3*. At seven days after LCMV Armstrong infection, PD-1^hi^ cells also had lower expression of effector-associated genes such as *Tbx21, Id2, Cxcr6 and Ccr5* and reduced levels of granzymes; in addition, these cells did not persist long term. In addition, during acute infection Tim-3^hi^ cells expressed higher levels of genes encoding effector-associated proteins (*Tbx21, Ifng, Gzmb)* and genes encoding exhaustion markers (*Pdcd1, Lag3, Tox).* However, unlike Tpex cells, our study reveals that Tim-3^hi^ effectors have better survival compared to Tim-3^lo^ CD8^+^ T cell effectors, as shown by active caspase-3 and Annexin V labelling *ex-vivo*. Furthermore, Tim-3^hi^ effectors have increased survival *in vitro*, which appears to be independent of Tim-3 ligands, based on antibody blocking (Fig. S4). Taken together, these data show that Tim-3^hi^ CD8 effectors are not only more differentiated, but they are also capable of better survival and may lead to better protection against viral infections.

## Acknowledgements

We thank the NIH Tetramer Core Facility (NIH Contract 75N93020D00005 and RRID:SCR_026557) for providing LCMV tetramers. We thank the Unified Flow Core for assistance with flow sorting and the Health Sciences Sequencing Core for planning and execution of RNA and TCR sequencing. This work was supported by NIH awards R01AI138504 and R01CA206517 to L.P.K., and by T32CA082084 to P. M.

## Notes

### Competing Interest Statement

The authors have declared no competing interest.

