## Supplemental Figures 1-4 for "Tim-3 Promotes Early Differentiation of Tbet^+^ Effector T Cells During Acute Viral Infection"

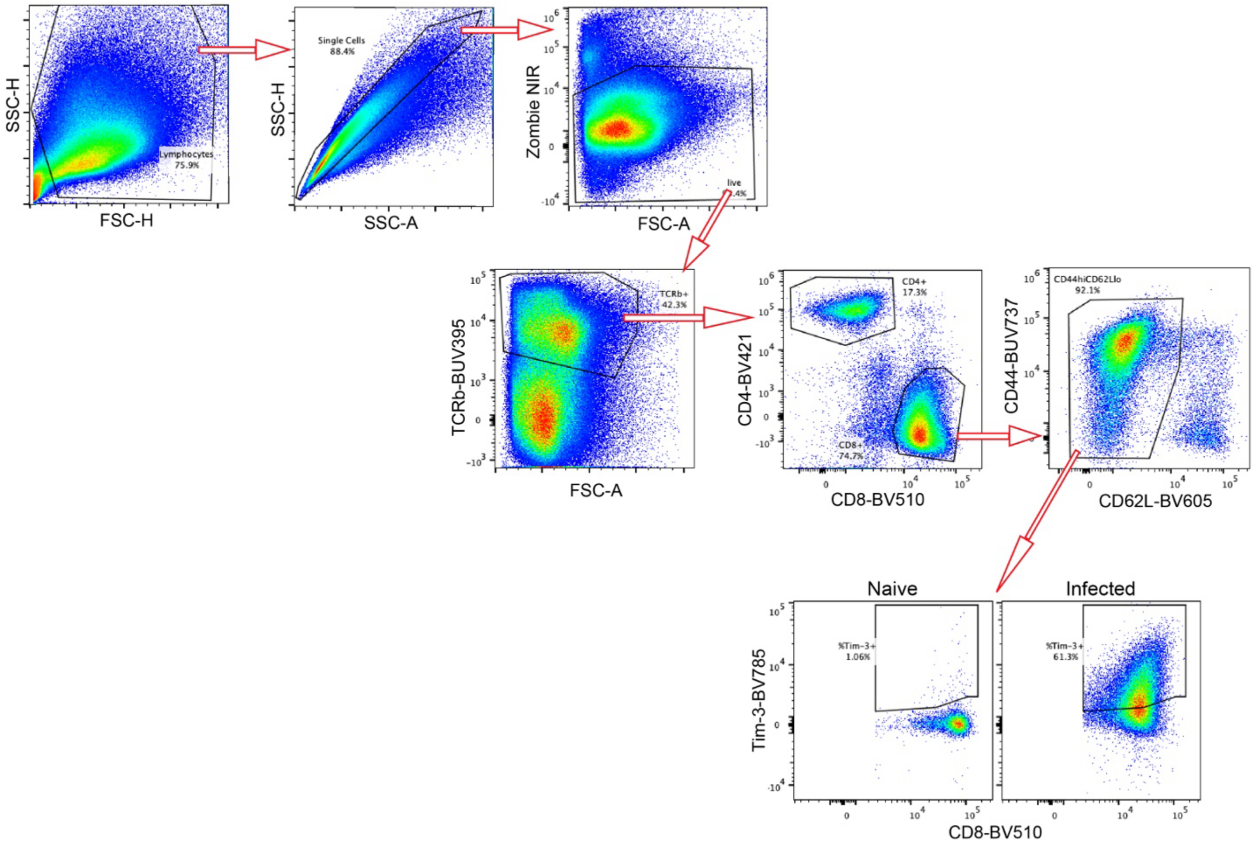


**Fig. S1: Representative flow cytometry gating strategy for analyzing antigen-experienced CD8+ T cells from the spleen of LCMV-infected mice.**

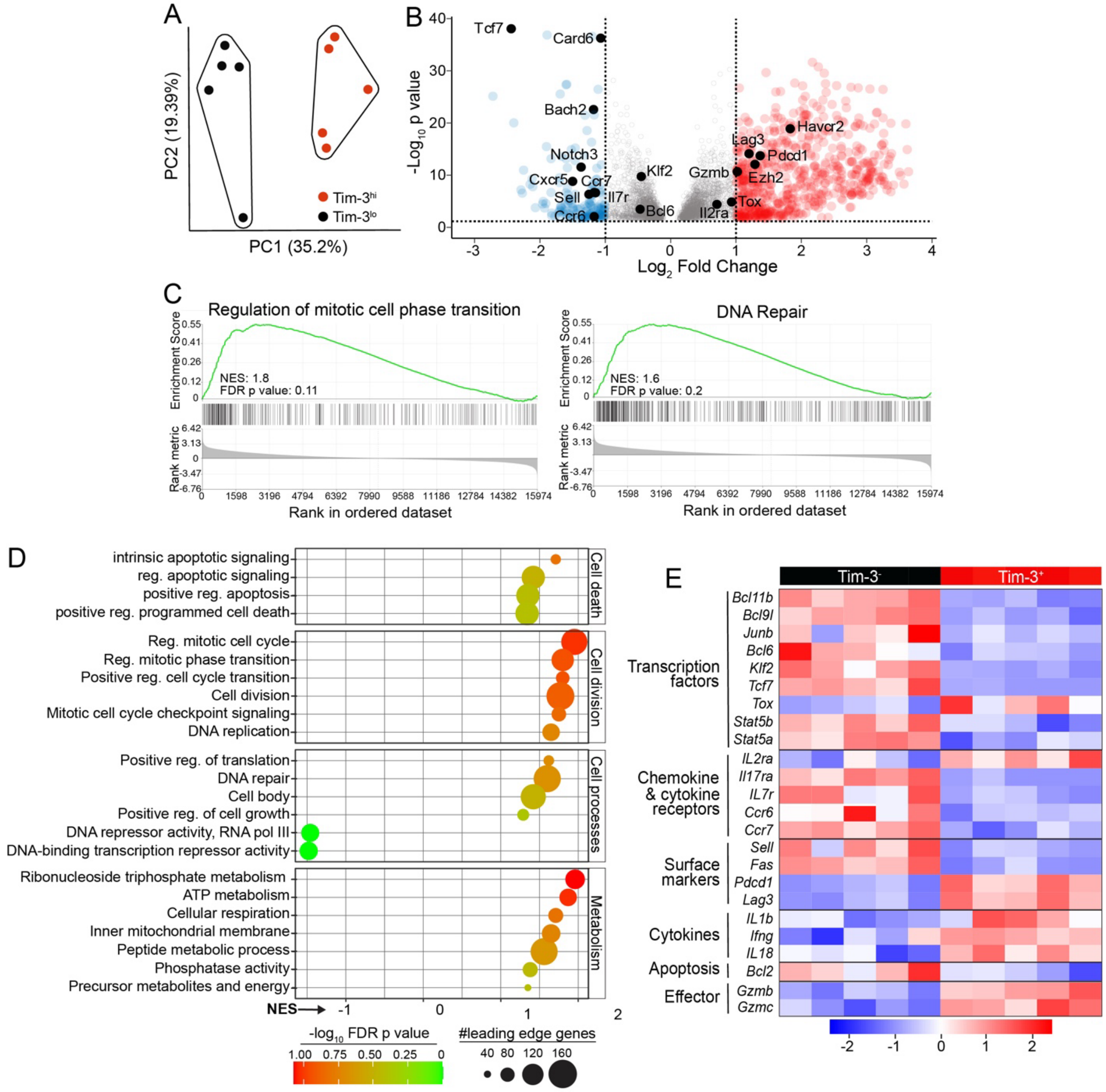


**Fig. S2: Upregulation of distinct pathways and effector genes in Tim-3$^{hi}$ CD8$^{+}$ effector T cells.** Bulk RNA sequencing was performed on Tim-3$^{lo}$ and Tim-3$^{hi}$ cells from C57BL/6 mice on day 8 after infection with LCMV Armstrong. CD44$^{hi}$CD62L$^{lo}$ CD8$^{+}$ T cells were sorted for sequencing. (**A**) Principal component analysis (PA). (**B**) Volcano plot depicting genes upregulated and downregulated in Tim-3$^{hi}$ vs. Tim-3$^{lo}$ CD8$^{+}$ T cells. (**C**) Selected GSEA plots, showing a positive coregulation of Tim-3 upregulation with pathways associated with cell cycle and DNA repair. (**D**) Dot plot representation of multiple GSEA pathways. A positive normalized enrichment score (NES) indicates upregulation of the indicated pathway in Tim-3$^{hi}$ T cells. (**E**) Heatmap showing the relative expression of selected effector-associated genes in individual samples.

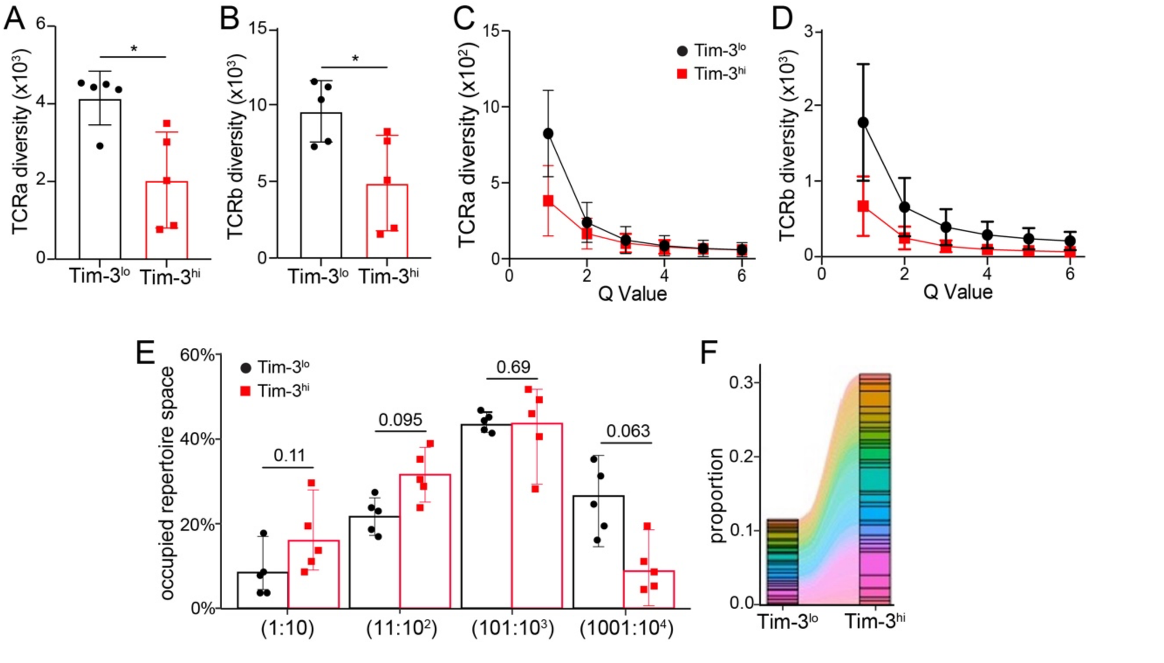


**Fig. S3: Focusing of the TCR repertoire in Tim-3$^{hi}$ CD8$^{+}$ effector T cells.** Bulk TCR sequencing was performed on the same samples described in Fig. S1. (**A-D**) Diversity of TCR$\alpha$ (**A,C**) and TCR$\beta$ (**B,D**) chains, derived with the Chao1 (**A-B**) and Hill (**C-D**) diversity estimators. **(E)** Occupied repertoire space of top 10, $10^2$, $10^3$ and $10^4$ clones. (**F**) Graph showing shared TCR$\beta$ clones between Tim-3$^{lo}$ and Tim-3$^{hi}$ CD8+ T cells. Each dot represents an individual mouse (n=5). Statistical significance was determined using Student's t-test. *p<0.05.

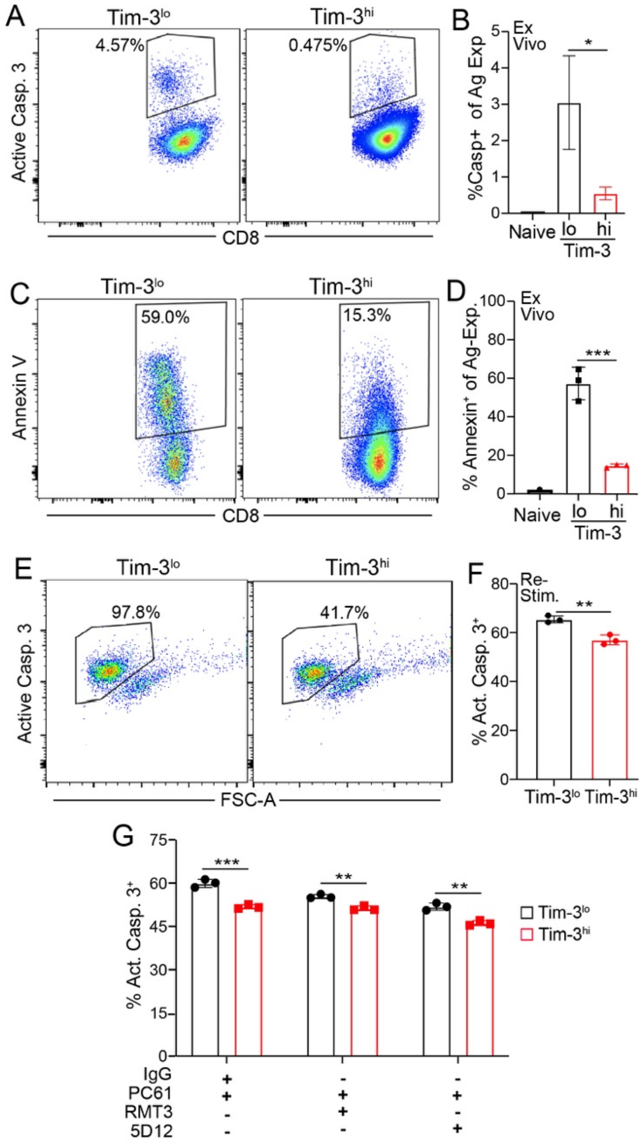


**Fig. S4: Tim-3$^{hi}$ effector T cells display enhanced survival after in vitro culture.** C57BL/6 mice were infected with 2x10$^5$ pfu of LCMV Armstrong. (**A-D**) Eight days later, splenic T cells were analyzed by flow cytometry for active caspase 3 (A-B) or surface PS, using Annexin V (C-D). (**E-G**) Tim-3$^{lo}$ and Tim-3$^{hi}$ CD8$^+$ T cells were sorted eight days after infection and cultured *in vitro* with 2μg/ml of plate bound anti-CD3 antibody in the absence of IL-2 for 24 hours. In panel G, the indicated antibodies were added to the culture. Each dot represents an individual animal (A-D) or technical replicates (G). Statistical significance was determined by student's t-test. *$p<0.05$, **$p<0.01$, ***$p<0.001$. Data are representative of two experiments.
